# Worldwide impacts of past and projected future land-use change on local species richness and the Biodiversity Intactness Index

**DOI:** 10.1101/311787

**Authors:** Samantha L.L. Hill, Ricardo Gonzalez, Katia Sanchez-Ortiz, Emma Caton, Felipe Espinoza, Tim Newbold, Jason Tylianakis, Jörn P. W. Scharlemann, Adriana De Palma, Andy Purvis

## Abstract

Although people have modified the world around us throughout human history, the ‘Great Acceleration’ has seen drivers such as land conversion, exploitation of natural populations, species introductions, pollution and human-induced climate change placing biodiversity under increasing pressure. In this paper we examine 1) how terrestrial species communities have been impacted over the last thousand years of human development and 2) how plausible futures defined by alternative socio-economic scenarios are expected to impact species communities in the future. We use the PREDICTS (Projecting Responses of Ecological Diversity In Changing Terrestrial Systems) database to model impacts of land-use change and human population on local species richness, community abundance, and biodiversity intactness using a mixed-effects modelling structure. Historical impacts are inferred through projection of model results onto maps of historical land use, provided by the land-use harmonization project, and gridded human population density (HYDE 3.1). Future impacts are explored using the Shared Socio-economic Pathway (SSP) scenarios. These scenarios detail five plausible global futures based upon socio-economic factors such as wealth, population, education, technology, and reliance on fossil fuels, and can be combined with Representative Concentration Pathway (RCP) scenarios to consider climate mitigation strategies. We project model results onto the gridded outputs of six SSP/RCP scenario combinations: SSP1/RCP2.6, SSP2/RCP4.5, SSP3/RCP7.0, SSP4/RCP3.4, SSP4/RCP6.0, and SSP5/RCP8.5. Historical trend lines show that most losses in local biodiversity are relatively recent, with 75% of all loss in both abundance-based Biodiversity Intactness Index and species richness occurring post-1800. Stark regional differences emerge in all future scenarios, with biodiversity in African regions undergoing greater losses than Oceania, North America and the European regions. Although climate change is expected to have severe detrimental impacts to biodiversity – which are not quantified in these results – it is important to consider how the climate change mitigation itself may also impact biodiversity. Our results suggest that strong climate change mitigation through biofuel production will detrimentally impact biodiversity: SSP4/RCP3.4 (with high biofuel mitigation) is predicted to see two times the decrease in abundance-based biodiversity intactness and three times the decrease in local species richness between 2015–2100 as is predicted for SSP4/RCP6.0 (with lower levels of mitigation). SSP4/RCP3.4 forecasts the greatest impact to average local species richness of all the SSP/RCP combinations with an average loss of 13% of local species richness projected to have occurred by 2100. SSP3/RCP7.0 – a scenario describing a globally segregated, and economically protectionist future with low climate change mitigation – has the worst impacts on abundance-based biodiversity intactness with an average loss of 26% of intactness by 2100. However, a brighter future is possible; SSP1/RCP2.6 describes a more sustainable future, where human populations are provided for without further jeopardising environmental integrity – in this scenario we project that biodiversity will recover somewhat, with gains in biodiversity intactness and species richness in many regions of the world by 2100.

## Introduction

Biodiversity indicators are important tools for supporting sustainability. However, most biodiversity indicators report only over a short time period leading up to the present day, because few underpinning observational time-series go back more than a few decades (Magurran et al. 2010). Furthermore, because indicators based on such time series do not embody a model of how biodiversity is affected by drivers, future projections must be based on simple extrapolation of recent trends (e.g., Tittensor et al. 2014), so it is not possible to compare biodiversity outcomes from alternative future pathways. Indicators that embody explicit links between drivers and biodiversity provide the potential for estimating how the state of the indicator has changed beyond the temporal range for which direct observations are available, enabling estimation not only of how the state of nature has changed up to now, but also how it will change in possible futures (e.g., Nicholson et al. 2012; IPBES Scenarios & Modelling assessment (IPBES 2016); Visconti et al. 2015; Purvis et al. 2018).

The PREDICTS modelling framework focuses on the biodiversity effects of land use and related pressure (Purvis et al. 2018), as these are still the dominant current pressures on terrestrial biodiversity worldwide (Foley et al. 2005; Sala et al. 2000). The statistical models linking biodiversity to drivers are underpinned by a large global and taxonomically broad database of terrestrial ecological communities facing land-use pressures (Hudson et al. 2014, 2017). The core assumption is that the relationships between the drivers and biodiversity estimated from these data remain constant over time (Purvis et al. 2018). Once these relationships are estimated statistically, the model coefficients are then crossed with global layers of the relevant drivers for any year of interest, to produce estimates of the desired biodiversity indicator or indicators.

The PREDICTS database was designed to be compatible with the harmonized land-use classes that Hurtt et al. (2011) used in their gridded historical maps of land use from 1500–2005 and their projections of land use from 2005–2100 under each of the four Representative Concentration Pathways (RCPS: van Vuuren et al. 2011) as implemented by a set of four Integrated Assessment Models (IAMs: Harfoot et al. 2014). This decision allowed Newbold et al. (2015) to estimate that land use and related pressures have reduced average local species-richness across the world’s terrestrial assemblages by 13.6%, with most of the decline concentrated in the 20th century. Newbold et al. (2015) also inferred that four different RCP x IAM combinations had very different implications for biodiversity by 2100: average species-richness was predicted to fall by a further 3.4% under MESSAGE 8.5 but to increase by 1.9% under MINICAM 4.5.

More recently, the PREDICTS framework has been extended to estimate Scholes & Biggs’ (2005) Biodiversity Intactness Index (BII: Newbold et al. 2016). BII is defined as the average abundance of a taxonomically and ecologically broad set of species in an area, relative to their abundances in an intact reference ecosystem (Scholes & Biggs 2005), and has been proposed as a potential indicator of whether global ecosystems are still within a ‘safe operating space’ in the Planetary Boundaries framework (Steffen et al. 2015). PREDICTS estimates BII by combining two statistical models; one of overall organismal abundance relative to an intact baseline, and one of compositional similarity to an intact baseline ecosystem (Newbold et al. 2016; Purvis et al. 2018). The global mean BII was estimated as being 84.6% (Newbold et al. 2016), which is below the proposed safe limit of 90% (Steffen et al. 2015), but no temporal trajectory of BII was estimated. Subsequently, De Palma et al. (2018) have combined statistical models for tropical and subtropical forest biomes with annual high-resolution estimates of land use to infer how BII changed from 2001–2012 in these biomes; but so far no global analysis has inferred temporal change in BII.

Here, we go beyond these previous analyses in two main ways. First, we apply the PREDICTS modelling framework for the first time to the five Shared Socioeconomic Pathways (SSPs: Riahi et al. 2017) developed as part of the sixth round of Intergovernmental Panel on Climate Change (IPCC) reports. This necessitated the re-curation of sites in the PREDICTS database to be compatible with the expanded set of land-use classes used by Hurtt et al. (in prep) in their new harmonization, LUH2. Second, we have improved the modelling of compositional similarity, enabling explanatory variables other than land use and distance to affect the compositional similarity between sites. We present estimates of how global average values of two indicators – species richness and abundance-based BII have changed between 850 and 2010, and their future trajectories to 2100 under each of six SSP/RCP combinations made available in the harmonized dataset.

## Methods

The PREDICTS database is a globally and taxonomically comprehensive database of site level measures of biodiversity within different gradients of human pressure (Hudson et al. 2017). Site-level data was extracted from the PREDICTS database in October 2017. The database contained 3.85 million rows of data, incorporating data on approximately 31 000 taxa from 32 000 sites and 767 studies.

Land use and land use intensity were assigned using data provided by the original source publications at the time of entry into the database. However, the Cropland, Plantation Forest and Pasture sites needed recuration to match the more detailed categories of the LUH2 projections. Timber plantations were removed from the PREDICTS Plantation Forest category, and the remaining Plantation Forest sites (i.e., those containing permanent woody crops such as fruit trees) were merged with Croplands. Managed Pasture and Rangelands were reassigned from the PREDICTS land use class of Pasture through information found in the site descriptions provided in the original primary literature sources as well as knowledge of broad regional patterns in the density of grazing (see Appendix). Information on the crop species under cultivation at the PREDICTS cropland sites was gleaned from the original primary literature source or from interview with the data provider. Crop species were classified as Annual or Perennial using the TRY database (Kattge et al. 2011). All crop species in the family Fabaceae were considered to be Nitrogen-fixing. Cropland sites were classified as Annual/Perennial/Nitrogen-fixing if the majority of crops identified at each site fell into one of these classes. Cropland sites that could not be classified in this manner were dropped from the analysis. For a description of the PREDICTS/LUH2 land use classes see supplemental information. Site level human population density was extracted from GRUMP (CIESIN 2011).

For each site, we calculated total abundance as the sum of all individuals of all species, and species richness as the number of present species. Abundance-based compositional similarity was calculated using the asymmetric Jaccard index (Chao et al. 2005) to provide a metric detailing the proportion of individuals in a comparison site that are of species that were also present in the baseline site. Studies that only sampled single species were excluded from the dataset. For abundance models, where sampling effort varied among sites within a study, abundance was first rescaled assuming that diversity increased linearly with sampling effort; such studies were excluded for models of compositional similarity. Abundance was also log transformed prior to modelling to improve the distribution; modelling with Poisson errors was not possible because, even prior to rescaling, many abundance values were not integers.

Local species richness and abundance-based compositional similarity models were run using a mixed-effects modelling framework, implemented in the R package lme4 (version 1.1–14 Bates et al. 2017). The random-effects structure was selected using Akaike’s Information Criterion (AIC) values (Zuur et al. 2009). Source, study and block level random intercepts were included within all models and a site level random intercept was employed within the species richness model to combat overdispersion. Random slopes were not considered because they led to problems with model convergence.

Explanatory variables included in the model selection process were human population density (ln(x+1) transformed), land use, land-use intensity, and a factor combining the two (which we term LUI). Study-level means of human population density and agricultural suitability were included as control variables, to avoid possible biases (e.g., sampling may be more thorough in studies conducted in more densely-populated regions). All continuous variables were modelled using linear relationships. The species richness model was modelled using a Poisson distribution and all other models used a Gaussian distribution (overall abundance having been log-transformed). The fixed effects model structure was selected using backwards stepwise selection based upon AIC values (Zuur et al. 2009).

The compositional similarity model followed the framework outlined in De Palma (2018). A matrix was constructed including all sites as rows and columns. Pairwise comparisons between all sites were calculated for 1) compositional similarity using the asymmetric Jaccard Index (Chao et al. 2005), 2) geographic distance (log transformed), 3) environmental distance (cube root transformed, selected over log transformation through analysis of residual variation), and 4) the difference in log human population density between sites. Only contrasts where the baseline site in the pairwise comparison contained Primary vegetation with Minimal use intensity were included within the model. Compositional similarity was transformed using logit transformation (to improve distribution of residuals, as compositional similarity is bounded between zero and one) and modelled against geographic distance, environmental distance, and the pairwise contrast of land uses as an interaction with human population density (of the second site in the matrix). The difference between human population density was also included as an additive effect to quantify the impact of the change from a pristine system with no human influence to a system with human influence as this is of interest as well as the absolute impact of human population.

Pressure maps for use in projections were derived as follows. Land use and human population density maps were obtained from the LUH2 dataset (Hurtt et al. in prep). The age of secondary vegetation was tracked from 800 using the transition rates from Hurtt et al. (in prep), and categorised as young secondary when age < 30 years, intermediate secondary when age >30 year and <50 years, and mature secondary when age > 50 years. A static use intensity map was produced using a model to predict likelihood of intensity through human population density and GlobCover (following Newbold et al. 2015).

Modelled coefficients were projected onto pressure maps to produce gridded maps of local species richness, total community abundance and abundance-based compositional similarity. The abundance maps was then multiplied by the compositional similarity map to produce maps of abundance-based BII (Newbold et al. 2016). Aggregated results for the globe and for IPBES subregions (Brooks et al. 2016) were calculated using arithmetic means weighted by net primary productivity for abundance results, and by vertebrate species richness for species-richness results, as in Newbold et al. (2015). Historic maps were produced from 800 to 2014 and future projections for each year from 2015 to 2100 were produced for the following SSP/RCP combinations: SSP1/RCP2.6, SSP2/RCP4.5, SSP3/RCP7.0, SSP4/RCP3.4, SSP4/RCP6.0, and SSP5/RCP8.5.

## Results

Historical trend lines reveal a dramatic decrease in average local biodiversity following the Industrial Revolution, with 75% of the reduction to date in both biodiversity intactness and local species richness occurring post-1800. However, there are regional variations. Biodiversity was impacted earlier within Central and Western Europe but then began to recover towards the end of the twentieth century (Figures 1 & 2). Conversely, regions such as the Caribbean, West Africa and South Asia experienced dramatic losses in biodiversity in the twentieth century (Figures 1 & 2).

**Figure 1.**
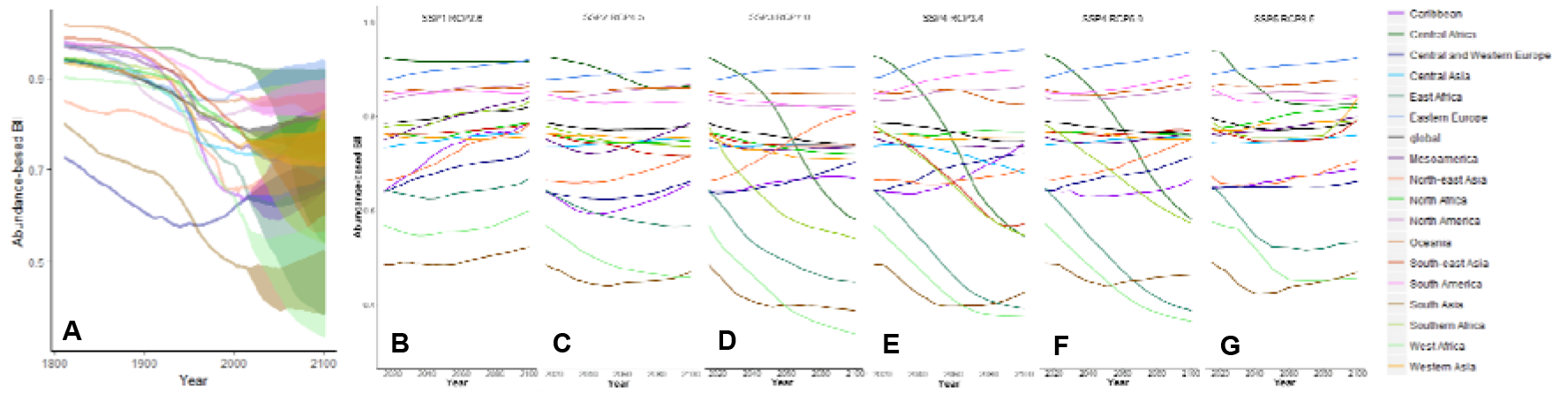
**Temporal trends in abundance-based BII at a global level and for each of the IPBES subregions. Trendlines show the average loss in BII from an unimpacted baseline. A) Trend from 1800 to 2100. Extent of variation between SSP/RCP future projections is indicated by shaded areas. B) to G). Global and regional trends for each SSP/RCP projection showing results for SSP1/RCP2.6, SSP2/RCP4.5, SSP3/RCP7.0, SSP4/RCP3.4, SSP4/RCP6.0, and SSP5/RCP8.5 respectively.**

**Figure 2.**
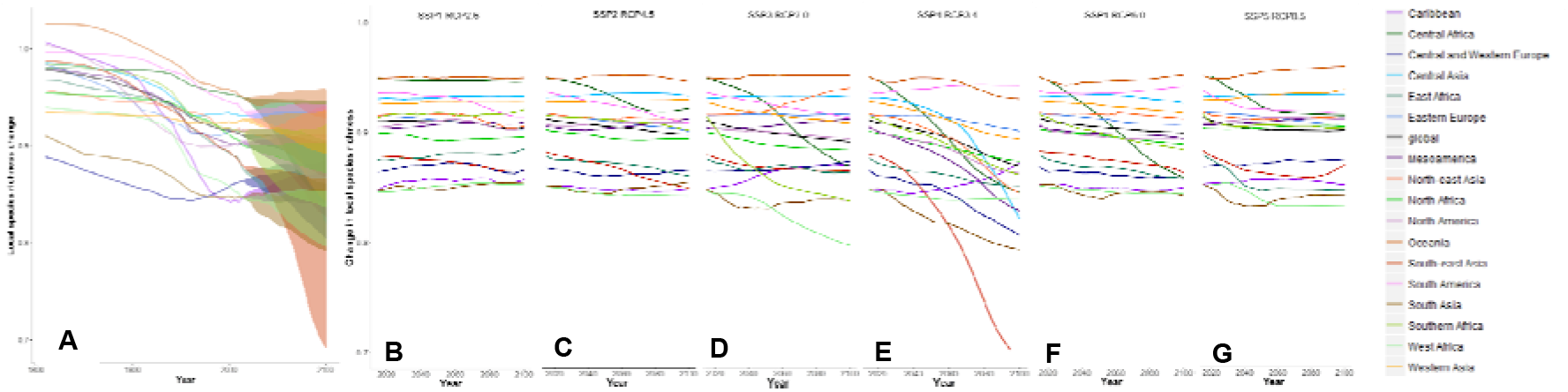
**Temporal trends in local species richness at a global level and for each of the IPBES subregions. Trendlines show the average loss in local species richness from an unimpacted baseline. A) Trend from 1800 to 2100. Extent of variation between SSP/RCP future projections indicated by shaded areas. B) to G). Global and regional trends for each SSP/RCP projection showing results for SSP1/RCP2.6, SSP2/RCP4.5, SSP3/RCP7.0, SSP4/RCP3.4, SSP4/RCP6.0, and SSP5/RCP8.5 respectively.**

Global mean abundance-based BII in 2015 is estimated to be 0.785 (Table 1), with all regions apart from Central Africa predicted to have a value of less than 0.90 (the proposed safe limit: Steffen et al. 2015). Global mean local species richness in 2015 is estimated to have been 0.901.

**Table 1.** **Average abundance-based BII for each SSP/RCP combination in 2015, 2050 and 2100.**

|  |  | SSP1/RCP2.6 | SSP2/RCP4.5 | SSP3/RCP7.0 | SSP4/RCP3.4 | SSP4/RCP6.0 | SSP5/RCP8.5 |
| --- | --- | --- | --- | --- | --- | --- | --- |
| Global | 2015 | 0.785 | 0.785 | 0.785 | 0.785 | 0.785 | 0.785 |
|  | 2050 | 0.799 | 0.770 | 0.760 | 0.770 | 0.769 | 0.761 |
|  | 2100 | 0.821 | 0.778 | 0.740 | 0.744 | 0.764 | 0.776 |
| Caribbean | 2015 | 0.641 | 0.641 | 0.641 | 0.641 | 0.641 | 0.641 |
|  | 2050 | 0.731 | 0.593 | 0.659 | 0.643 | 0.627 | 0.653 |
|  | 2100 | 0.786 | 0.656 | 0.669 | 0.739 | 0.661 | 0.681 |
| Central Africa | 2015 | 0.924 | 0.924 | 0.924 | 0.924 | 0.924 | 0.924 |
|  | 2050 | 0.917 | 0.895 | 0.834 | 0.807 | 0.819 | 0.826 |
|  | 2100 | 0.917 | 0.863 | 0.580 | 0.544 | 0.578 | 0.815 |
| Central and Western Europe | 2015 | 0.639 | 0.639 | 0.639 | 0.639 | 0.639 | 0.639 |
|  | 2050 | 0.685 | 0.622 | 0.651 | 0.683 | 0.669 | 0.641 |
|  | 2100 | 0.728 | 0.661 | 0.703 | 0.717 | 0.709 | 0.654 |
| Central Asia | 2015 | 0.734 | 0.733 | 0.733 | 0.734 | 0.734 | 0.734 |
|  | 2050 | 0.750 | 0.741 | 0.739 | 0.727 | 0.744 | 0.736 |
|  | 2100 | 0.765 | 0.752 | 0.735 | 0.677 | 0.748 | 0.750 |
| East Africa | 2015 | 0.645 | 0.644 | 0.643 | 0.643 | 0.643 | 0.645 |
|  | 2050 | 0.630 | 0.588 | 0.510 | 0.502 | 0.507 | 0.518 |
|  | 2100 | 0.668 | 0.567 | 0.448 | 0.392 | 0.386 | 0.527 |
| Eastern Europe | 2015 | 0.877 | 0.877 | 0.876 | 0.877 | 0.877 | 0.877 |
|  | 2050 | 0.898 | 0.886 | 0.897 | 0.918 | 0.902 | 0.892 |
|  | 2100 | 0.920 | 0.900 | 0.905 | 0.935 | 0.931 | 0.911 |
| Mesoamerica | 2015 | 0.751 | 0.751 | 0.750 | 0.751 | 0.751 | 0.752 |
|  | 2050 | 0.793 | 0.723 | 0.729 | 0.731 | 0.738 | 0.761 |
|  | **2100** | 0.841 | 0.784 | 0.734 | 0.734 | 0.758 | 0.788 |
| **North Africa** | **2015** | 0.764 | 0.764 | 0.764 | 0.764 | 0.764 | 0.764 |
|  | **2050** | 0.756 | 0.744 | 0.738 | 0.756 | 0.759 | 0.785 |
|  | **2100** | 0.786 | 0.748 | 0.720 | 0.765 | 0.755 | 0.807 |
| **North America** | **2015** | 0.834 | 0.835 | 0.835 | 0.835 | 0.835 | 0.834 |
|  | **2050** | 0.856 | 0.847 | 0.824 | 0.847 | 0.833 | 0.838 |
|  | **2100** | 0.872 | 0.866 | 0.824 | 0.859 | 0.857 | 0.834 |
| **South America** | **2015** | 0.846 | 0.846 | 0.846 | 0.846 | 0.846 | 0.846 |
|  | **2050** | 0.85 | 0.826 | 0.829 | 0.874 | 0.850 | 0.820 |
|  | **2100** | 0.851 | 0.830 | 0.812 | 0.895 | 0.882 | 0.830 |
| **North-east Asia** | **2015** | 0.663 | 0.663 | 0.663 | 0.663 | 0.663 | 0.663 |
|  | **2050** | 0.710 | 0.660 | 0.721 | 0.653 | 0.681 | 0.649 |
|  | **2100** | 0.782 | 0.715 | 0.807 | 0.686 | 0.747 | 0.697 |
| **Oceania** | **2015** | 0.852 | 0.852 | 0.852 | 0.852 | 0.852 | 0.852 |
|  | **2050** | 0.853 | 0.852 | 0.847 | 0.851 | 0.840 | 0.852 |
|  | **2100** | 0.863 | 0.859 | 0.848 | 0.823 | 0.865 | 0.865 |
| **South Asia** | **2015** | 0.485 | 0.485 | 0.484 | 0.485 | 0.485 | 0.485 |
|  | **2050** | 0.484 | 0.440 | 0.397 | 0.401 | 0.436 | 0.421 |
|  | **2100** | 0.522 | 0.470 | 0.388 | 0.426 | 0.462 | 0.465 |
| **Southern Africa** | **2015** | 0.778 | 0.777 | 0.777 | 0.777 | 0.777 | 0.778 |
|  | **2050** | 0.795 | 0.747 | 0.617 | 0.687 | 0.690 | 0.746 |
|  | **2100** | 0.831 | 0.735 | 0.541 | 0.546 | 0.570 | 0.781 |
| **West Africa** | **2015** | 0.570 | 0.569 | 0.568 | 0.568 | 0.568 | 0.570 |
|  | **2050** | 0.555 | 0.489 | 0.430 | 0.442 | 0.451 | 0.473 |
|  | **2100** | 0.599 | 0.458 | 0.338 | 0.375 | 0.364 | 0.452 |
| **South-east Asia** | **2015** | 0.764 | 0.764 | 0.763 | 0.764 | 0.764 | 0.764 |
|  | **2050** | 0.766 | 0.739 | 0.739 | 0.690 | 0.753 | 0.737 |
|  | **2100** | 0.784 | 0.714 | 0.738 | 0.569 | 0.765 | 0.783 |
| **Western Asia** | **2015** | 0.762 | 0.762 | 0.761 | 0.761 | 0.761 | 0.762 |
|  | **2050** | 0.754 | 0.757 | 0.724 | 0.748 | 0.757 | 0.774 |
|  | **2100** | 0.757 | 0.755 | 0.708 | 0.749 | 0.746 | 0.828 |

The SSP/RCP combinations allow the examination of how differing socioeconomic scenarios will impact biodiversity at global and regional scales. SSP1/RCP2.6 shows both an overall global positive response in biodiversity intactness and broadly consistent positive responses within regions (Figure 1B). For all other scenarios the regional results are less consistent. For instance, when examining the projections for SSP3/RCP7.0 and SSP5/RCP8.5, most African regions show declines in both biodiversity intactness and local species richness, whereas some Asian and European regions are predicted to see an overall improvement in biodiversity (Figures 1D, 1G, 2D & 2G).

The comparison of SSP4/RCP3.4 and SSP4/RCP6.0 allows the evaluation of the impact to biodiversity from land use changes aimed at mitigating global temperature increases (note that our framework does not assess the impacts of temperature increase themselves). This mitigation has negative impacts, causing a three times greater decrease in local species richness, and two times greater decrease in abundance-based biodiversity intactness, from 2015–2100 for SSP4/RCP3.4 compared to that for SSP4/RCP6.0.

The global map of present-day abundance-based biodiversity intactness (here illustrated through SSP3/RCP7.0) shows relatively low values throughout much of Western Europe and Eastern North America; however, the lowest levels are seen in areas where high population density overlaps with high land conversion, for instance, much of India and Northern China (Figure 3). A comparison of abundance-based BII in 2015 and 2050 reveals increases over much of Western Europe but declines over much of Central and Southern Africa (Figure 4).

**Figure 3.**
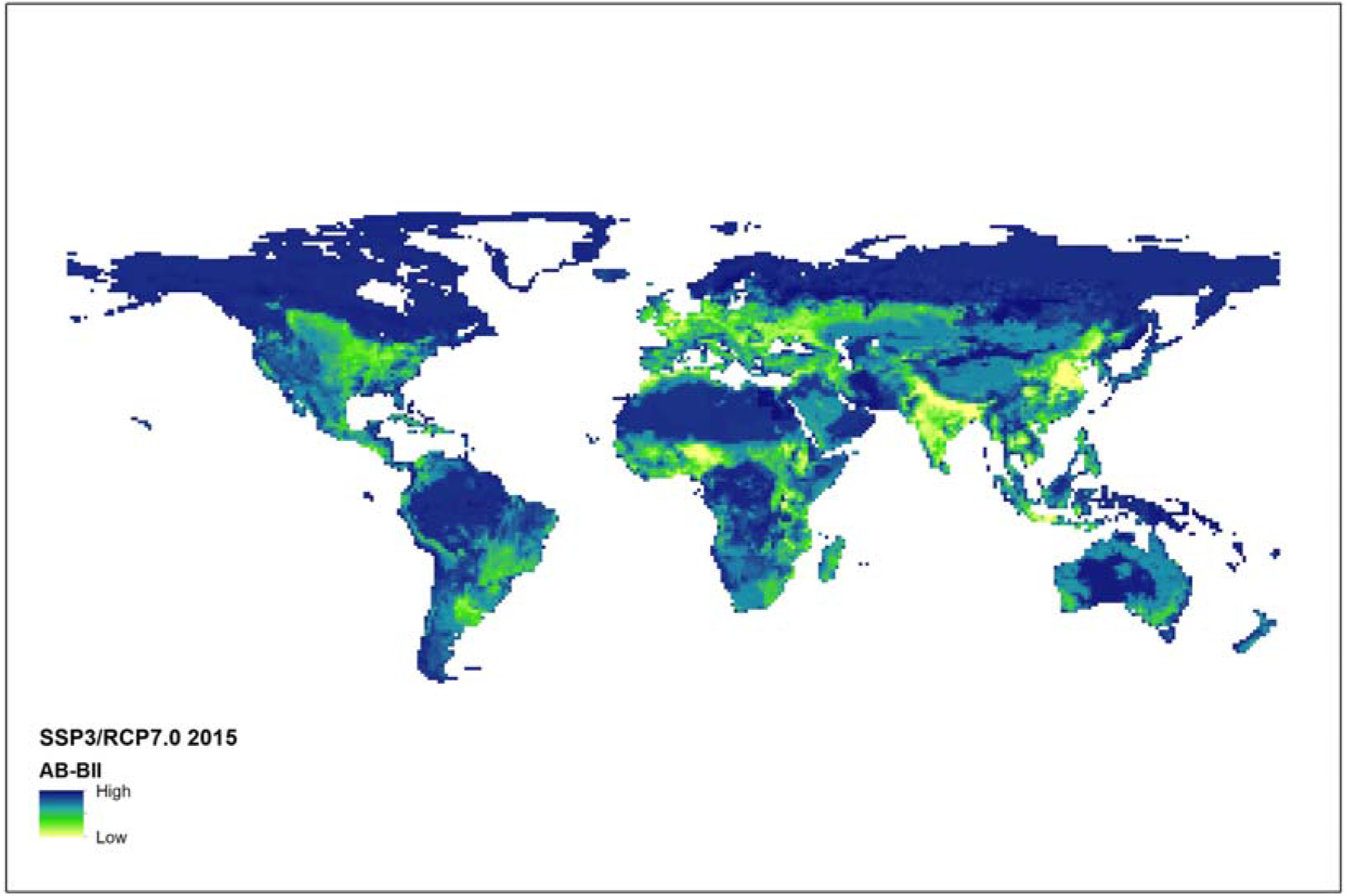
**Abundance-based biodiversity intactness projected for 2015 under SSP3/RCP7.0.**

**Figure 4.**
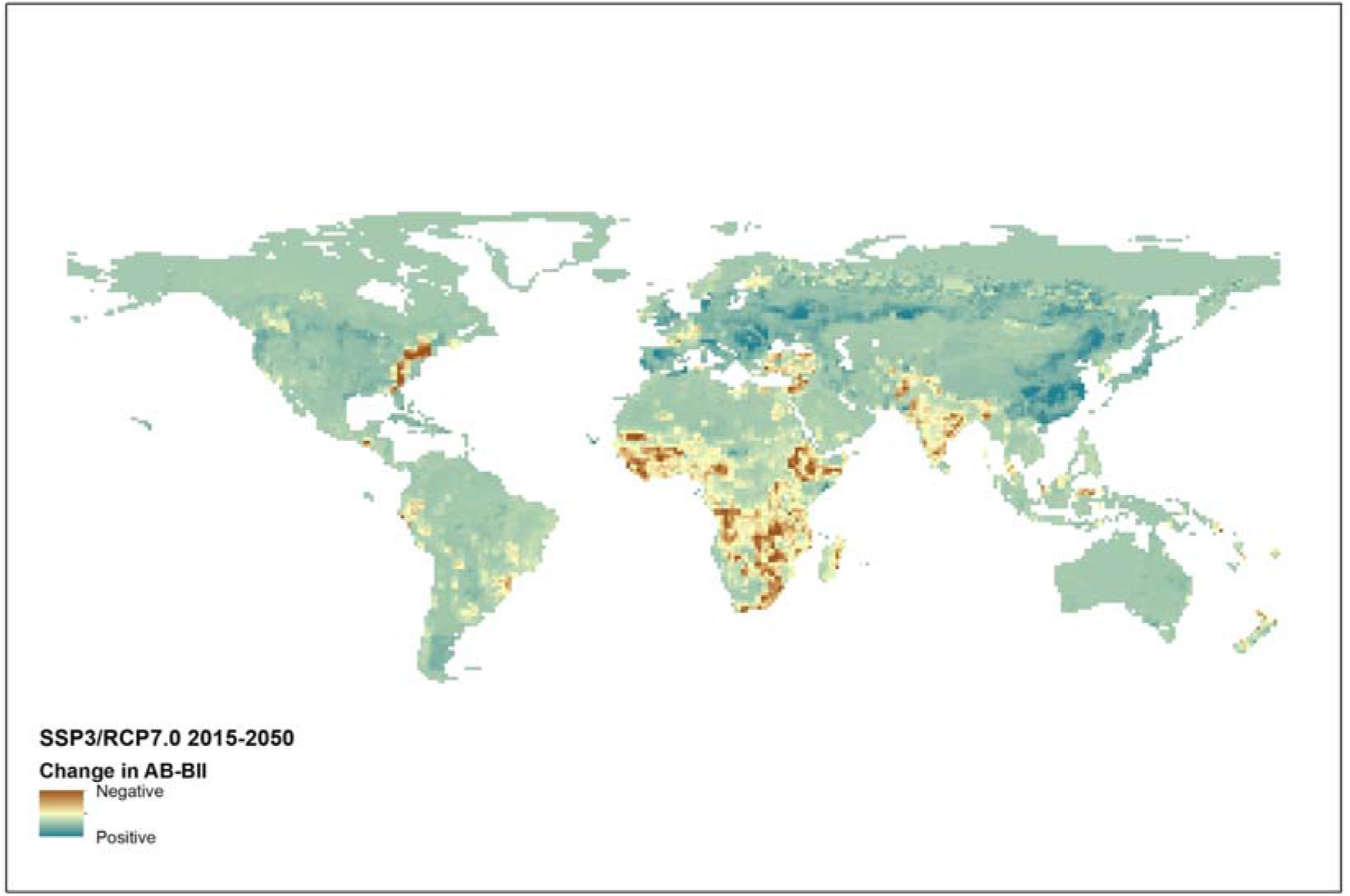
**Change in abundance-based BII projected to occur between 2015 and 2050 by SSP3/RCP7.0.**

## Discussion

Our projections of the future for biodiversity under varying plausible socio-economic scenarios reveal stark regional differences with African regions faring worse than European and Asian regions under five of the six SSP/RCP combinations. Only SSP1/RCP2.6 showed overall improvement in biodiversity globally and in most regions. This optimistic scenario describes a sustainable future, broadly in line with the United Nations Sustainable Development Goals (SDGs), where population growth is minimised, global levels of education are improved, and agricultural demand is minimised through sustainable practices and behaviour (Riahi et al. 2017).

Our global estimate of mean abundance-based BII, at 0.785, is somewhat lower than the 0.846 estimated by Newbold et al. (2016). Several factors probably contribute to this difference. The major factor is likely to be that our models of compositional similarity are able to incorporate more pressures than did those of Newbold et al. (2016) because we use the full set of pairwise comparisons rather than using only a subset. The resulting models include significant negative effects of human population density and land-use intensity on compositional similarity - variables omitted from the models of Newbold et al. (2016). Purvis et al. (2018) discuss some remaining reasons why our modelling approach may still be overestimating BII; the most important in the context of this paper is that our models do not consider biotic effects of climate change or other drivers that have a different spatial pattern from land use and human population density.

Our map of present-day abundance-based BII shows geographic differences from the map presented by Newbold et al. (2016). Most obviously, we infer a markedly higher level of BII across much of Australia (compare Figure 3 with Newbold et al.’s Figure S4) than Newbold et al. (2016) who found surprisingly low values of BII. This difference is the clearest example of where the refined land-use classes of LUH2 are an improvement over the classes in the original land-use harmonization (Hurtt et al. 2011); much of Australia is currently Rangeland, with a much more intact biota (higher abundance, richness and compositional similarity to baseline sites) than Managed Pasture, but these two land-use classes were united as Pasture by Hurtt et al. (2011) and in Newbold et al.’s (2016) models. The refinement of Hurtt et al.’s (2011) single, broad Cropland class into five LUH2 classes presented challenges for data curation and modelling, and would require a larger database to model fully, but also provides more nuanced models that better reflect - albeit still coarsely - different effects on local biodiversity.

There are some difficulties in interpreting differences among projections for the different SSP/RCP combinations considered here. Most SSPs are explored at only a single level of radiative forcing (i.e., at a single RCP) within the LUH2 data, and most SSP/RCP combinations were originally developed within a single Integrated Assessment Model (Riahi et al. 2017). Additionally, the harmonized LUH2 data represent in some sense a compromise among the various IAMs, none of which resolve land use into exactly the same land-use classes used here. All three of these difficulties arise from sensible practical decisions taken in the face of resource constraints, but could potentially be overcome with further work. Extending LUH2’s coverage of SSP x RCP combinations within and among IAMs, though it cannot lead to a full-matrix design (because, for instance, not all RCPs can be achieved under all SSPs) would permit much more sophisticated exploration of the differences among projections. Aligning PREDICTS’ sites directly to the land-use classes within each IAM would reduce the potential for harmonization to cause - or to remove - differences among scenarios.

